# Composition is the core driver of the language-selective network

**DOI:** 10.1101/436204

**Authors:** Francis Mollica, Evgeniia Diachek, Zachary Mineroff, Hope Kean, Matthew Siegelman, Steven T. Piantadosi, Richard Futrell, Peng Qian, Evelina Fedorenko

## Abstract

The fronto-temporal language network responds robustly and selectively to sentences. But the features of linguistic input that drive this response and the computations these language areas support remain debated. Two key features of sentences are typically confounded in natural linguistic input: words in sentences a) are semantically and syntactically combinable into phrase- and clause-level meanings, and b) occur in an order licensed by the language’s grammar. Inspired by recent psycholinguistic work establishing that language processing is robust to word order violations, we hypothesized that the core linguistic computation is composition, and, thus, can take place even when the word order violates the grammatical constraints of the language. This hypothesis predicts that a linguistic string should elicit a sentence-level response in the language network as long as the words in that string can enter into dependency relationships as in typical sentences. We tested this prediction across two fMRI experiments (total N=47) by introducing a varying number of local word swaps into naturalistic sentences, leading to progressively less syntactically well-formed strings. Critically, local dependency relationships were preserved because combinable words remained close to each other. As predicted, word order degradation did not decrease the magnitude of the BOLD response in the language network, except when combinable words were so far apart that composition among nearby words was highly unlikely. This finding demonstrates that composition is robust to word order violations, and that the language regions respond as strongly as they do to naturalistic linguistic input as long as composition can take place.

---

A left-lateralized network of anatomically and functionally inter-connected brain regions selectively supports language processing (e.g., Fedorenko, Behr, & Kanwisher, 2011). The regions of this “language network” respond to both i) word meanings, and ii) combinatorial semantic/syntactic processing (e.g., Keller, Carpenter, & Just, 2001; Fedorenko, Hsieh, Nieto-Castañón, Whitfield-Gabrieli, & Kanwisher, 2010; Fedorenko, Nieto-Castanon, & Kanwisher, 2012; Bautista & Wilson, 2016). The magnitude of neural responses in these regions, as measured with diverse brain imaging techniques, appears to scale with how language-like the input is, with strongest responses elicited by sentences, and progressively lower responses elicited by phrases, lists of unconnected words, pseudowords, and foreign/indecipherable speech (e.g., Fedorenko et al., 2010; Bedny, Pascual-Leone, Dodell-Feder, Fedorenko, & Saxe, 2011; Pallier, Devauchelle, & Dehaene, 2011; Vagharchakian, Dehaene-Lambertz, Pallier, & Dehaene, 2012; Fedorenko et al., 2016; Scott, Gallée, & Fedorenko, 2017; Hultéen, Schoffelen, Uddén, Lam, & Hagoort, 2019). But what features of the linguistic stimulus and what associated linguistic computations drive the language network’s response? In particular, sentences – its preferred stimulus – both a) contain word pairs that are semantically and syntactically combinable into phrases and clauses, and b) have the word order constrained by the rules of the language. Here we evaluate a hypothesis that the core linguistic computation has to do with combining words into phrases and clauses, and that this computation does not depend on word order (i.e., can take place even when the word order is not licensed by the language’s grammar). A key prediction of this hypothesis is that a linguistic string should elicit a sentence-level response in the language network as long as the words in that string are combinable.

The motivation for this hypothesis is two-fold. First, all languages reflect the structure of the world (e.g., Mikolov, Sutskever, Chen, Corrado, & Dean, 2013; Pennington, Socher, & Manning, 2014), including both broad generalizations (e.g., that properties can apply to objects or entities, that entities can engage in actions, or that some actions can affect objects) and particular contingencies (e.g., which specific properties apply to which objects, which specific entities engage in which actions, etc.). This knowledge of the world along with lexical knowledge (knowledge of word meanings) determines which words in the linguistic input are combined to form phrases and clauses during language comprehension. For example, the words *tasty* (a property, denoted by an adjective) and *apple* (an object, denoted by a noun) are combinable into a phrase, in this case one with a plausible meaning, but the words *tasty* and *ate* (an action, denoted by a past tense verb) cannot be combined because adjectives are not typically dependents of verbs like *taste*. In contrast, although many accounts of syntactic representation and processing have emphasized word order as a key cue to building syntactic structures (e.g., Bever, 1970; Kimball, 1973), languages across the world vary widely in the rigidity of their word order constraints, with many languages exhibiting highly flexible orderings, pointing to a more limited role of word order at least in those languages (e.g., K. Hale, 1983; Dryer & Haspelmath, 2013; Jackendoff & Wittenberg, 2014). As a result, combinability of words into phrases and clauses, but not strict word order, appears to be a universal feature of linguistic input that our language processing mechanisms must be able to handle.

And second, recent work in psycholinguistics has shown that our sentence interpretation mechanisms are well designed for coping with errors – including morpho-syntactic agreement errors and word swaps – as long as a plausible meaning is recoverable (e.g., Ferreira, Bailey, & Ferraro, 2002; Levy, 2008b; Levy, Bicknell, Slattery, & Rayner, 2009; Gibson, Bergen, & Piantadosi, 2013; Traxler, 2014). These coping mechanisms are pervasive enough to interfere with our ability to detect errors during proofreading (e.g., Schotter, Bicknell, Howard, Levy, & Rayner, 2014) and to make grammaticality judgments for sentences with easily correctable syntactic errors compared to clearly grammatical/ungrammatical sentences (Mirault, Snell, & Grainger, 2018). As a result, if the core linguistic computation implemented in the language-selective cortex has to do with combining words into phrases and clauses, form-based errors may be irrelevant as long as they do not impede this process.

To test this hypothesis, we used a novel manipulation to examine neural responses to sentences where word order is degraded (to varying extents), but local dependency relationships are preserved. In particular, naturalistic sentences were gradually degraded by increasing the number of local word swaps (Figure 1), which broke syntactic dependencies and led to progressively less syntactically well-formed strings (Table 1). Critically, local semantic and syntactic relationships were preserved. The degree of local combinability can be formally estimated using tools from information theory (Shannon & Weaver, 1963). Naturalistic linguistic input is characterized by relatively high pointwise mutual information (PMI) among words within a local linguistic context, and it falls off for word pairs spanning longer distances (e.g., Li, 1990; Lin & Tegmark, 2017; Futrell, Qian, Gibson, Fedorenko, & Blank, 2019). Our local-word-swap manipulation maintained approximately the same level of local mutual information as that observed in typical linguistic input. As can be seen in Figure 2e, the conditions with 1, 3, 5, and even 7 word swaps (Scr1, Scr3, Scr5, Scr7) have similar local PMI levels to the intact condition (see Methods for details). To evaluate the importance of locality for building dependency relationships, in one condition (in Experiment 2), we scrambled words within each sentence in a way so as to minimize local PMI and thus break local inter-word relationships. In this condition, local PMI is comparable to that of a list of unconnected words (see ScrLowPMI and Word-list conditions in Figure 2e). Participants read these materials – presented one word at a time – while undergoing fMRI, and blood oxygenation level dependent (BOLD) responses were examined in language-selective regions defined using a separate localizer task (Fedorenko et al., 2010, Figure 2a).

**Figure 1:**
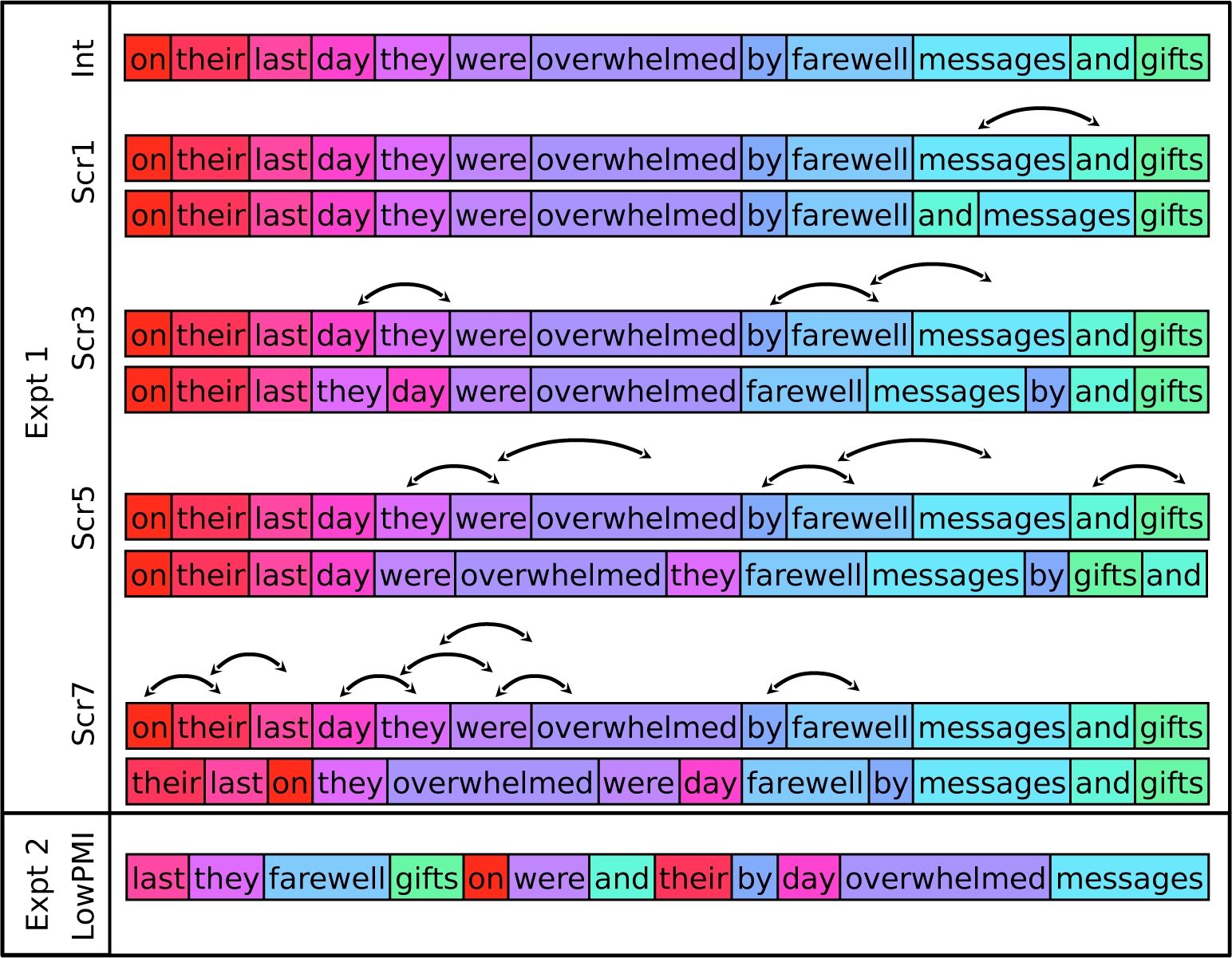
A sample item from the critical experiment; colors are used to illustrate the increasing degradedness (i.e., the color spectrum becomes progressively more discontinuous with more swaps). a. The schematic of the procedure used to create the scrambled-sentence conditions in Experiment 1. b. A sample stimulus from the ScrLowPMI condition in Experiment 2.

**Figure 2:**
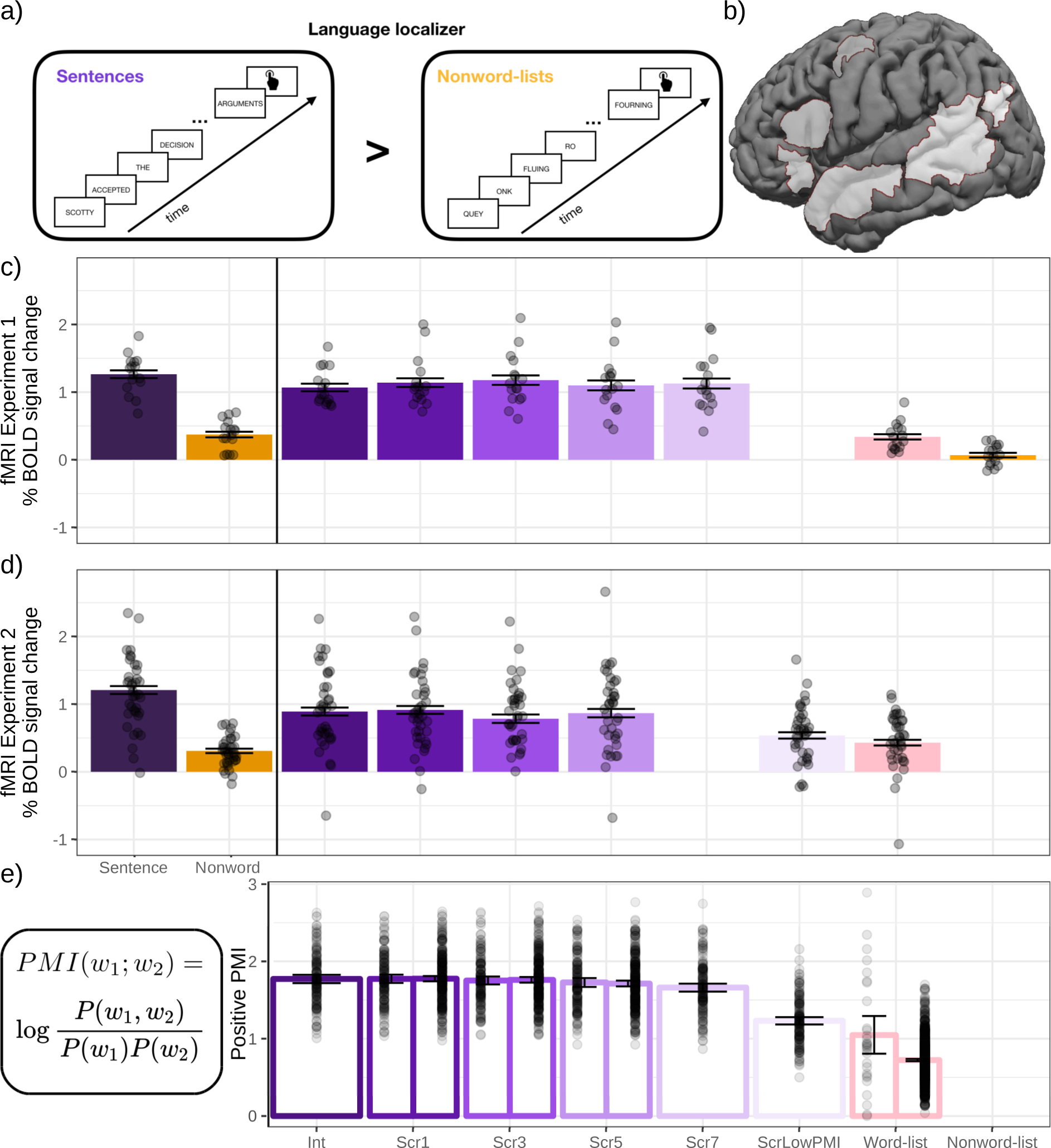
a. The schematic of the language localizer task used to define the language-responsive areas. b. The parcels used to define the language-responsive areas. In each participant, the top 10% of most localizer-responsive voxels within each parcel were taken as that participant’s region of interest. Replicating prior work (Fedorenko et al., 2010), the localizer effect—estimated using across-runs cross-validation to ensure independence—was highly robust in both experiments (ps¡0.0001). c-d. Neural responses (in % BOLD signal change relative to fixation) to the conditions of the language localizer and Experiments 1 (n=16) and 2 (n=32). e. The formula for computing pointwise mutual information (PMI) (see Materials and Methods for details), and average positive PMI values for the materials in Experiments 1 and 2 (N.B.: Slightly different scramblings of the materials for the Scr1, Scr3, and Scr5 conditions were used in the two experiments; hence two bars (left=Experiment 1) for each of these conditions.)

**Table 1:**
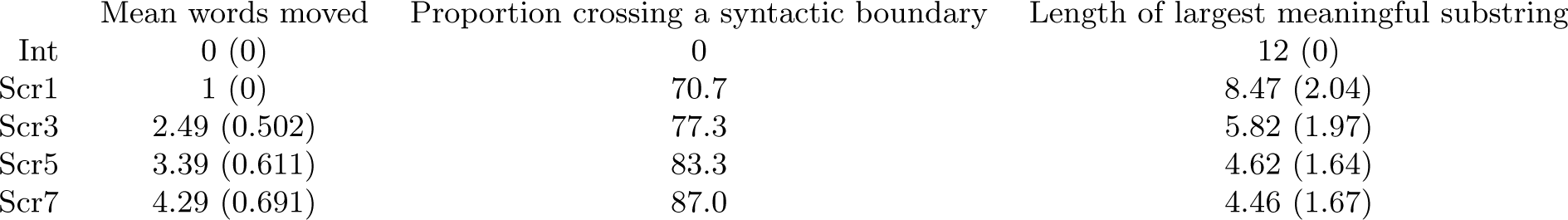
Description of the stimuli in Experiment 1 and 2a. Mean (SD)

|  | Mean words moved | Proportion crossing a syntactic boundary | Length of largest meaningful substring |
| --- | --- | --- | --- |
| Int | 0 (0) | 0 | 12 (0) |
| Scr1 | 1 (0) | 70.7 | 8.47 (2.04) |
| Scr3 | 2.49 (0.502) | 77.3 | 5.82 (1.97) |
| Scr5 | 3.39 (0.611) | 83.3 | 4.62 (1.64) |
| Scr7 | 4.29 (0.691) | 87.0 | 4.46 (1.67) |

If the core function of the language processing mechanisms is to combine words into phrases and clauses, and this process is robust to word order violations, we would expect the neural response to remain high as long as local PMI is similar to that observed in naturalistic linguistic input, but to drop for the condition where local PMI is low. If, on the other hand, composition critically depends on word order, such that it is hindered or altogether blocked in cases where the word order violates the grammatical rules of the language, or if the core linguistic computation has to do with word-order-based parsing, then we would expect the neural response to decrease as the word order becomes more degraded. It is also possible, on this hypothesis, that there would be a non-linearity in the response across conditions, with an increase for conditions with a small number of word swaps, which are relatively easily correctable with the cost carried by the language areas, and then a drop for conditions with a larger number of swaps.

To foreshadow the key results, we found that the fMRI BOLD response in the language areas does not decrease, relative to the response to its preferred stimulus (sentences), as long as mutual information among nearby words remains as high as in typical linguistic input, allowing for composition. However, scrambling a sentence so as to minimize local mutual information, blocking composition, leads to the response dropping to the level of that for a list of unconnected words. These results support the idea that composition is the core computation implemented in the language network, and this computation is robust to word order violations.

## Methods

### Participants

Forty-seven individuals (age 18−48, average age 22.8; 31 females) participated for payment (Experiment 1: *n* = 16; Experiment 2: *n* = 32; one individual participated in both Experiment 1 and Experiment 2, and one individual participated in Experiment 2 twice, once in version a and once in version b, as described below, for a total of 49 scanning sessions across the 47 participants; for the participant who participated in Experiment 2 twice, the data were combined across the two sessions). (We included twice as many participants in Experiment 2 to ensure that the critical result in Experiment 1 was not due to insufficient power.) Forty-one participants were right-handed, as determined by the Edinburgh handedness inventory (Oldfield, 1971), or by self-report; the remaining six left-handed/ambidextrous individuals showed typical left-lateralized language activations in the language localizer task (see Willems, Van der Haegen, Fisher, & Francks, 2014, for arguments to include left-handers in cognitive neuroscience research). All participants were native speakers of English from the Boston community. Four additional participants were scanned (for Experiment 2) but excluded from the analyses due to excessive head motion or sleepiness, and/or failure to perform the behavioral task. All participants gave written informed consent in accordance with the requirements of MIT’s Committee on the Use of Humans as Experimental Subjects.

### Experimental Design and Materials

In both experiments, each participant completed a) a version of the language localizer task (Figure 2a Fedorenko et al., 2010), which was used to identify language-responsive areas at the individual-subject level, and b) the critical sentence comprehension task (in 30/49 scanning sessions, participants completed the localizer task in the same session as the critical task, for the remaining 19 sessions, the localizer came from an earlier session; see Mahowald and Fedorenko (2016), for evidence of the stability of the localizer activation maps across sessions). In addition, each participant completed a spatial working memory task (Fedorenko et al., 2011), used in some control analyses to characterize brain regions sensitive to sentence scrambling, as described below. Some participants further completed one or two additional tasks for unrelated studies. The language localizer task was always completed first; the order of all other tasks varied across participants. The entire scanning session lasted approximately 2 hours.

### Language localizer

Participants passively read sentences and lists of pronounceable nonwords in a blocked design. The *Sentences > Nonwords* contrast targets brain regions sensitive to high-level linguistic processing (Fedorenko et al., 2010). The robustness of this contrast to materials, modality of presentation, language, and task has been previously established (Fedorenko et al., 2010; Fedorenko, 2014; Scott et al., 2017). In this version of the localizer, the sentences were constructed to vary in content and structures used, and the nonwords were created using the Wuggy software (Keuleers & Brysbaert, 2010), to match the phonotactic properties of the nonwords to those of the words used in the Sentence condition. Each trial started with 100 ms pre-trial fixation, followed by a 12-word-long sentence or a list of 12 nonwords presented on the screen one word/nonword at a time at the rate of 450 ms per word/nonword. Then, a line drawing of a hand pressing a button appeared for 400 ms, and participants were instructed to press a button whenever they saw this icon, and finally a blank screen was shown for 100 ms, for a total trial duration of 6 s. The simple button-press task was included to help participants stay awake and focused. Each block consisted of 3 trials and lasted 18 s. Each run consisted of 16 experimental blocks (8 per condition), and five fixation blocks (14 s each), for a total duration of 358 s (5 min 58 s). Each participant performed two runs. Condition order was counterbalanced across runs.

### Spatial working memory task (used in some control analyses)

Participants had to keep track of four (easy condition) or eight (hard condition) locations in a 3×4 grid (Fedorenko et al., 2011). In both conditions, participants performed a two-alternative, forced-choice task at the end of each trial to indicate the set of locations that they just saw. The *Hard > Easy* contrast targets brain regions sensitive to general executive demands (e.g., Duncan & Owen, 2000; Duncan, 2010). Fedorenko, Duncan, and Kanwisher (2013) have shown that the regions activated by this task are also activated by a wide range of tasks that contrast a difficult vs. an easier condition Hugdahl, Raichle, Mitra, and Specht (2015, see also). Each trial lasted 8 s (see Fedorenko et al., 2011, for details). Each block consisted of 4 trials and lasted 32 s. Each run consisted of 12 experimental blocks (6 per condition), and 4 fixation blocks (16 s each), for a total duration of 448 s (7 min 28 s). Forty-five participants performed two runs; the remaining two participants performed one run. Condition order was counterbalanced across runs when participants performed two runs.

### Critical task in Experiment 1

#### Design and materials

Participants read sentences with correct word order (Intact (Int)) and sentences with progressively more scrambled word orders created by an increasing number (between 1 and 7) of local word swaps (Scrambled (Scr) 1, 3, 5, and 7; Figure 1), as well as two control conditions: lists of unconnected words and lists of nonwords. At the end of each trial, participants were presented with a word (in the sentence and word-list conditions) or a nonword (in the nonword-list condition) and asked to decide whether this word/nonword appeared in the preceding trial.

To create the sentence materials, we extracted 150 12-word-long sentences from the British National corpus (Burnard, 2000). We then permuted the word order in each sentence via local swaps, to create the scrambled conditions. In particular, a word was chosen at random and switched with one of its immediate neighbors. This process was repeated a specified number of times. Because one random swap can directly undo a previous swap, we ensured that the manipulation was successful by calculating the edit distance. (The code used to create the scrambled conditions is available at OSF: https://osf.io/y28fz/.) We chose versions with 1, 3, 5, and 7 swaps in order to i) limit the number of sentence conditions to five, while, at the same time, ii) covering a range of degradedness levels. The materials thus consisted of 150 sentences with five versions each (Int, Scr1, Scr3, Scr5, and Scr7), for a total of 750 strings. These were distributed across five experimental lists following a Latin Square design, so that each list contained only one version of a sentence and 30 trials of each of the five conditions. Any given participant saw the materials from just one experimental list, and each list was seen by 2-4 participants.

To characterize the sentence materials in greater detail, as critical for interpretation (see Discussion), we performed three analyses on the materials used in Experiments 1 and 2a (Table 1). First, we manually annotated the number of words that were moved in each scrambled condition (where a move is defined as a rightward or leftward movement of a word across one or more words). For example, if *the dog chased the cat* was scrambled by 3 swaps to *dog the cat chased the*, two words (*the* and *cat*) have moved; and if it was scrambled by 3 swaps to *the chased the cat dog*, only one word (*dog*) has moved. As expected, this value increased gradually from the least to the most scrambled condition (i.e., from 1 in the Scr1 condition to 4.29 in the Scr7 condition), suggesting that there were more opportunities to break syntactic dependencies as the number of swaps increased. Second, we manually annotated the stimuli for the proportion of swaps that crossed a constituent boundary in the original sentence. This number increased gradually from 70.7% in the Scr1 condition to 87% in the Scr7 condition. This analysis ensures that the scrambling procedure broke syntactic dependencies, even in the condition with a single swap, and did not simply swap words within constituents. And finally, we computed the length of the largest contiguous grammatical and meaningful substring (whether or not that substring was present in the original sentence). This value decreased gradually from 8.47 words in the Scr1 condition to 4.46 words in the Scr7 condition.

The word-list condition consisted of sequences of 12 real words (173 unique words: 55.5% nouns, 15.6% verbs, 22.5% adjectives, and 6.4% adverbs; average word length: 7.19 phonemes, standard deviation 1.43 (Weide, 1998); average log frequency: 1.73, standard deviation 0.80 (Brysbaert, New, & Keuleers, 2012)), and the nonword-list condition consisted of sequences of 12 nonwords (there were actually four different nonword-list conditions – a manipulation not of interest to the current study; we averaged the responses across the four nonword-list conditions in the analyses). (The nonwords used in this experiment were generated differently from the nonwords used in the language localizer task. In particular, they were created from real words by introducing some number of letter replacements keeping local phonotactics intact. We do not make any direct comparisons between nonword conditions across experiments, so this difference is of no consequence.) The word-list and nonword-list materials were the same across participants. (All the materials are available at OSF: https://osf.io/y28fz/)

### Computing mutual information values

To estimate the likelihood of dependencies among nearby words, we used pointwise mutual information (PMI), a metric from information theory (Fano, 1961; Church & Hanks, 1990), which measures the mutual dependence between variables (in this case, words). Positive PMI values suggest a dependence between words based on their overlap in contexts of use. Negative and near-zero PMI values suggest the absence of a dependence. Following word2vec (Mikolov et al., 2013), we used a sliding four-word window to extract local word pairs from each 12-word string. This is equivalent to collecting the bigrams, 1-skip-grams, and 2-skip-grams from each string.

For each word pair, we calculated PMI as follows:

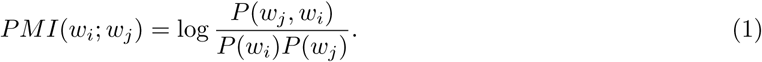

Probabilities were estimated using the Google N-gram corpus (Michel et al., 2010) and ZS (Smith, n.d.) with Laplace smoothing (*α* = 0.1). For each 12-word string, we averaged across the positive PMI values for all word pairs occurring within a four-word sliding window. (The code for computing PMI is available at OSF: https://osf.io/y28fz/.) Although PMI encompasses both semantic and syntactic dependence, it down-weighs the contribution of high frequency, closed-class words, like determiners, pronouns, and prepositions, given that it reflects inter-word association beyond the simple frequency of co-occurrence. As can be seen in Figure 2e, local PMI across the four scrambled conditions (Scr1, Scr 3, Scr5, and Scr 7) is as high as that in the intact (Int) condition. (Given that the sentences in the intact condition were drawn from a corpus, their local PMI values likely reflect average local PMI in typical linguistic input.) This operationalization is a coarse measure that collapses over finer-grained distinctions that may affect the formation of semantic and syntactic dependencies (e.g. and reviews, Bemis & Pylkkänen, 2011; Pylkkänen, Bemis, & Elorrieta, 2014; Pylkkänen, 2016, 2019). However, to the extent that this operationalization can account for patterns of neural (in this case, BOLD) responses and thus yield insights about the workings of the language system, it holds theoretical and empirical value.

### Procedure

Participants read sentences, scrambled sentences, word lists, and nonword lists in an event-related fMRI design. Each trial lasted 8 s and consisted of the presentation of the stimulus (a sequence of 12 words/nonwords presented one at a time in the center of the screen with no punctuation, for 500 ms each, in black capital letters on a white background), followed by a blank screen for 300 ms, followed by a memory probe presented in blue font for 1,200 ms, followed again by a blank screen for 500 ms. The memory probe came from the preceding stimulus on half of the trials. For the sentences, the probes were uniformly distributed across the beginning (first four words), middle (middle four words), or end (last four words) of the sentence; for the word and nonword lists, the probes were uniformly distributed across the 12 positions. Incorrect probes were the shuffled correct probes from other sequences in the same condition.

The trials in each experimental list (300 total; 30 trials per condition, where the conditions included the intact sentence condition, four scrambled sentence conditions, the word-list condition, and four nonword-list conditions) were divided into six subsets corresponding to six runs. Each run lasted 480 s (8 min) and consisted of 8 s * 50 trials (5 per condition) and 80 s of fixation. The optseq2 algorithm (Dale, 1999) was used to create condition orderings and to distribute fixation among the trials so as to optimize our ability to de-convolve responses to the different conditions. Condition order varied across runs and participants. Most participants (n = 13) performed 5 runs; the remaining three participants performed 4 or 3 runs due to time constraints.

### Critical task in Experiment 2

#### Design and materials

Experiment 2 was designed both 1) to assess the robustness of the results in Experiment 1, in line with increasing emphasis on replicability in cognitive neuroscience (e.g., Poldrack et al., 2017; Siegelman, Blank, Mineroff, & Fedorenko, 2019; Uddén et al., 2019), and 2) to directly evaluate the locality constraint on semantic composition. In particular, as discussed above, in typical linguistic input, semantic and syntactic dependencies tend to be local (e.g., Futrell, Mahowald, & Gibson, 2015). As a result, our linguistic processing mechanisms are plausibly optimized for building complex meanings within local linguistic contexts. For example, if words *tasty* and *apple* occur within the same sentence, but are separated by eight other words, we may be less likely to combine them compared to cases where they occur in close proximity to one another. To evaluate the importance of locality for the engagement of the composition mechanisms, we included a manipulation where words were scrambled within a sentence in a way that minimizes local PMI. If locality is important, this condition should elicit a lower neural response compared to the conditions with high local PMI because participants would not be engaging in composition.

As in Experiment 1, participants read sentences with correct word order (Int) and sentences with progressively more scrambled word orders (Scr 1, 3, and 5). The materials for these scrambled conditions were identical to those in Experiment 1 for half of the participants, and different permutations of the same intact stimuli for the other half. Because, as expected, the results were almost identical across these two versions of the materials, we report the results for all participants together.

The condition with 7 word swaps (Scr7) was replaced by a condition where each pair of nearby content words was separated as much as possible within the 12-word string, so as to minimize local mutual information (Figure 1). We focused on separating nearby content words because those carry the most information in the signal (Shannon & Weaver, 1963) and contribute to positive PMI values as noted above. Take, for example, one of our intact sentences: *Larger firms and international companies tended to offer the biggest pay rises*. First, the content words were given a fixed order that maximized the sum of the distances between adjacent content words (two content words are considered adjacent in the original string if they have no content words between them): e.g., *larger international tended biggest rises firms companies offer pay*. This process was repeated for the function words (e.g., *and the to*). Then, the ordered function words were embedded in the center of the ordered content words (i.e., *larger international tended biggest rises and the to firms companies offer pay*), which maximizes the distances between adjacent content words in the original sentence. (The code is available at OSF: https://osf.io/y28fz/.) The manipulation was effective, leading to a significant drop in local mutual information (Figure 2e). If locality is important for building inter-word relationships, then minimizing the likelihood of dependency formation within local contexts should lead to a drop in the neural response, similar to what is observed during the processing of unconnected word-lists (e.g., Fedorenko et al., 2010; Pallier et al., 2011).

In addition to the five sentence conditions, we included five word-list conditions that were matched in terms of their lexical properties word-for-word to the sentence conditions. In particular, each of 876 unique words in the sentence conditions was replaced by a different word of the same syntactic category (using the following set: nouns, verbs, adjectives, adverbs, and closed-class words), similar in length (+/-0.03 phonemes, on average (Weide, 1998)) and frequency (+/-0.23 log (lf), on average (Brysbaert et al., 2012)). (Due to a script error, 11 words got replaced by the same word as the original word, and 6 words got replaced by a word of a different part of speech.) We included the same number of word-list conditions as sentence conditions to match the distribution of sentence and word-/nonword-list conditions in Experiment 1. However, in the analyses, we averaged the responses across the five word-list conditions given that there is no reason to expect differences among them.

The materials were distributed across five experimental lists; any given participant saw the materials from just one list, and each list was seen by 5-7 participants. As in Experiment 1, at the end of each trial, participants were presented with a word and asked to decide whether this word appeared in the preceding trial (see Results for behavioral performance).

### Procedure

The procedure was identical to that in Experiment 1 except that the memory probe was uniformly distributed across the 12 positions in every condition. Most participants (n=30) performed 5 or 6 runs; the remaining two participants performed 4 or 3 runs due to time constraints.

### fMRI data acquisition

Structural and functional data were collected on the whole-body 3 Tesla Siemens Trio scanner with a 32-channel head coil at the Athinoula A. Martinos Imaging Center at the McGovern Institute for Brain Research at MIT. T1-weighted structural images were collected in 179 sagittal slices with 1 mm isotropic voxels (TR = 2,530 ms, TE = 3.48 ms). Functional, blood oxygenation level dependent (BOLD) data were acquired using an EPI sequence (with a 90° flip angle and using GRAPPA with an acceleration factor of 2), with the following acquisition parameters: thirty-one 4 mm thick near-axial slices, acquired in an interleaved order with a 10% distance factor; 2.1 mm×2.1 mm in-plane resolution; field of view of 200 mm in the phase encoding anterior to posterior (A *>* P) direction; matrix size of 96 mm x 96 mm; TR of 2,000 ms; and TE of 30 ms. Prospective acquisition correction (Thesen, Heid, Mueller, & Schad, 2000) was used to adjust the positions of the gradients based on the participant’s motion one TR back. The first 10 s of each run were excluded to allow for steady-state magnetization.

### fMRI data preprocessing and first-level analysis

First-level analyses were conducted in SPM5 (we used an older version of the software here due to the use of these data in other projects spanning many years and hundreds of subjects); critical second-level analyses were performed using custom MATLAB and R scripts. Each subject’s data were motion corrected (realignment to the mean image using second-degree b-spline interpolation) and normalized into a common brain space, the Montreal Neurological Institute (MNI) template (normalization was estimated for the mean image using trilinear interpolation) and resampled into 2 mm isotropic voxels. The data were then smoothed with a 4 mm Gaussian filter and high-pass filtered (at 200 s). The task effects in both the language localizer task and the critical experiment were estimated using a General Linear Model (GLM) in which each experimental condition was modeled with a boxcar function (corresponding to a block or event) convolved with the canonical hemodynamic response function (HRF). The model also included first-order temporal derivatives of these effects, as well as nuisance regressors representing entire experimental runs and offline-estimated motion parameters.

### Language fROI definition and response estimation

For each participant, functional regions of interest (fROIs) were defined using the Group-constrained Subject-Specific (GSS) approach (Fedorenko et al., 2010; Julian, Fedorenko, Webster, & Kanwisher, 2012), whereby a set of parcels or “search spaces” (i.e., brain areas within which most individuals in prior studies showed activity for the localizer contrast) is combined with each individual participant’s activation map for the same contrast. To define the language fROIs, we used six parcels (Figure 2b) derived from a group-level representation of data for the *Sentences > Nonwords* contrast in 220 participants (a set of participants scanned in our lab). These parcels included three regions in the left frontal cortex: two located in the inferior frontal gyrus (LIFG and LIFGorb), and one located in the middle frontal gyrus (LMFG); and three regions in the left temporal and parietal cortices spanning the entire extent of the lateral temporal lobe and extending into the angular gyrus (LAntTemp, LPostTemp, and LAngG). (These parcels were similar to the parcels reported originally in Fedorenko et al. (2010), except that the two anterior temporal regions were collapsed together, and the two posterior temporal regions were collapsed together.) Following much prior work in our group, individual fROIs were defined by selecting—within each parcel—the top 10% of most localizer-responsive voxels based on the *t*-values for the *Sentences > Nonwords* contrast. Responses (in percent BOLD signal change units) to the relevant critical experiment’s conditions, relative to the fixation baseline, were then estimated in these fROIs. So, the input to the critical statistical analyses consisted of—for each participant—a value (percent BOLD signal change) for each of 10 conditions in each of the six language fROIs. Further, for Experiment 1, responses were averaged across the four nonword-list conditions, leaving a total of seven conditions; and for Experiment 2, responses were averaged across the five word-list conditions, leaving a total of six conditions. In the critical analyses (Figure 2c-d and Table 5), we consider the language network as a whole (treating regions as random effects; see below) given the abundant evidence that the regions of this network form an anatomically (e.g., Saur et al., 2008; Axer, Klingner, & Prescher, 2013) and functionally integrated system, as evidenced by strong inter-regional correlations during rest and language comprehension (e.g., I. Blank, Kanwisher, & Fedorenko, 2014; Paunov, Blank, & Fedorenko, 2019) and by correlations in effect sizes across the regions (Mineroff, Blank, Mahowald, & Fedorenko, 2018), but we additionally report the six language fROIs’ individual profiles and associated statistics (Figure 3 and Table 6). (In addition, to facilitate comparisons with other datasets, we include individual participants’ whole-brain contrast maps for all the individual conditions relative to the fixation baseline on OSF: https://osf.io/y28fz/.)

**Figure 3:**
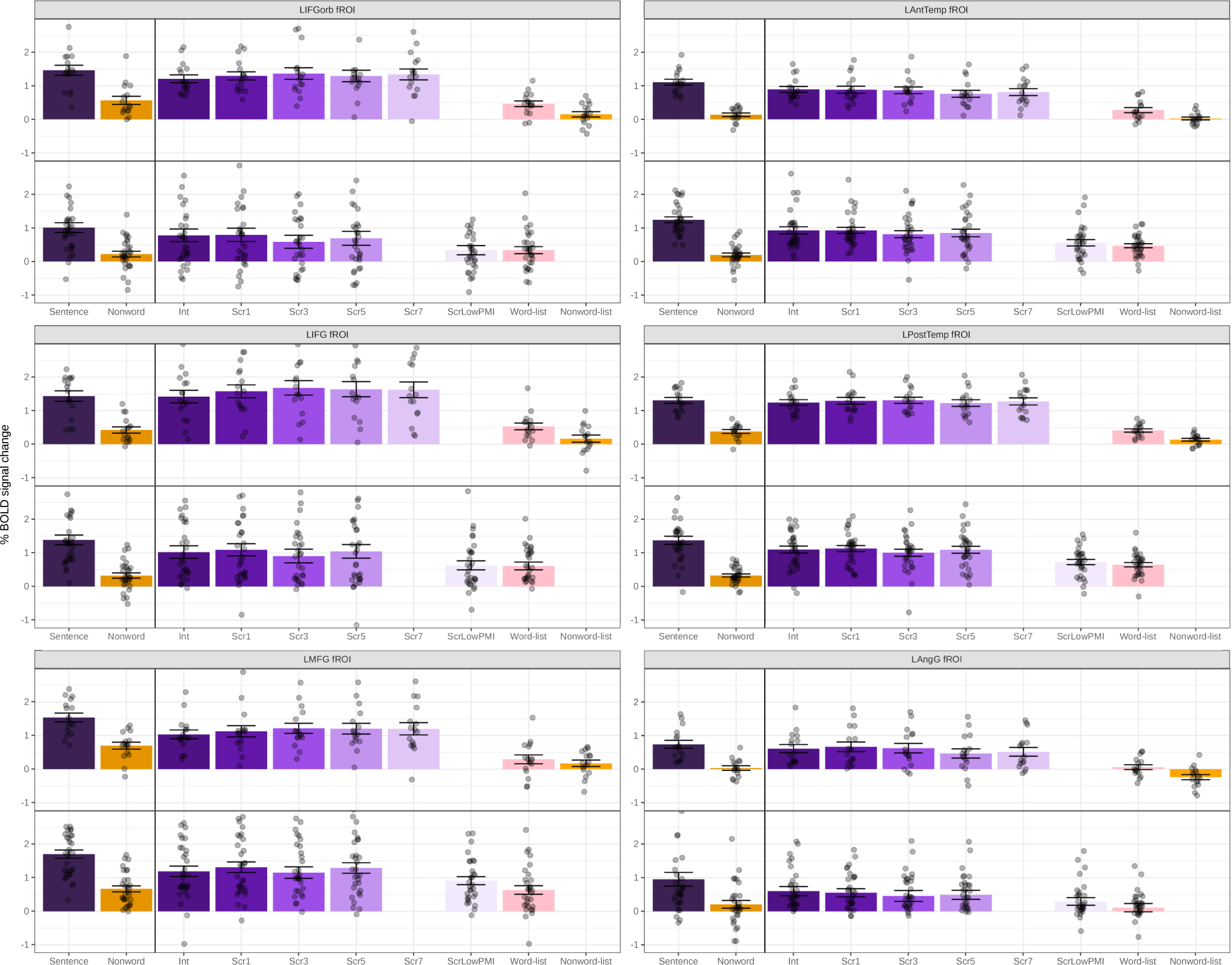
Neural responses (in % BOLD signal change relative to fixation) to the conditions of the language localizer and Experiments 1 (top panel) and 2 (bottom panel) in each of the six language fROIs.

### Statistical tests

To compare the average change in BOLD response across conditions, we conducted a mixed effect linear regression model with maximal random effect structure (Barr, Levy, Scheepers, & Tily, 2013), predicting the level of response with a fixed effect and random slopes for Condition, and random effects for Region of Interest (ROI) and Participant. To further compare the average change in BOLD response across conditions in each ROI separately, we conducted a mixed effect linear regression model with maximal random effect structure, predicting the level of response with a fixed effect and random slopes for Condition, and random effects for Participant. Condition was dummy-coded with Intact sentences as the reference level. Models were fit separately for Experiment 1 and Experiment 2 using the brms package (Buärkner et al., 2017) in R (Team, 2017) to interface with Stan (Stan Development Team, 2018).

### Behavioral naturalness rating study

To ensure that our scrambling manipulation was successful (in that human comprehenders would show sensitivity to it in some behavioral measure), 76 participants recruited through Amazon.com’s Mechanical Turk rated the naturalness of the sentence stimuli used in Experiment 1 on a 7-point scale (from 1=unnatural to 7=natural). On each trial, participants were presented with a single stimulus on the screen along with the scale. The endpoints of the scale were labeled. Participants responded by selecting a discrete point on the scale and then pressing the “Enter” key on their keyboard to move to the next trial. As in the fMRI study, the materials were distributed across five experimental lists (150 trials each) following a Latin Square design. Each list contained only one version of a sentence and 30 trials of each of the five conditions (Int, Scr1, Scr3, Scr5, Scr7). Any given participant saw the materials from just one experimental list. Due to a computer error, one list was administered to 16 participants; other lists were seen by 15 participants each.

The ratings were analyzed using a mixed effect linear regression model with a fixed effect and random slopes for Condition, and random effects for Participant and Item. To demonstrate the effectiveness of the manipulation at every level, Condition was backwards difference coded. As can be seen in Figure 4a and Table 2, every increase in degradation was associated with a significant decrease in perceived naturalness, although with diminishing returns. Thus participants were robustly sensitive to the scrambling manipulation. (The presentation code and data are available at OSF: https://osf.io/y28fz/.)

**Table 2:** The results of a mixed effect linear regression for the acceptability rating data. * denotes significant difference.

|  | Estimate | Est. Error | 95% Credible Interval |  |
| --- | --- | --- | --- | --- |
| Grand Mean | 3.54* | 0.08 | 3.38 | 3.69 |
| Scr1 - Int | -2.04* | 0.11 | -2.25 | -1.83 |
| Scr3 - Scr1 | -1.42* | 0.08 | -1.57 | -1.26 |
| Scr5 - Scr3 | -0.56* | 0.05 | -0.67 | -0.46 |
| Scr7 - Scr5 | -0.23* | 0.04 | -0.30 | -0.15 |

**Figure 4:**
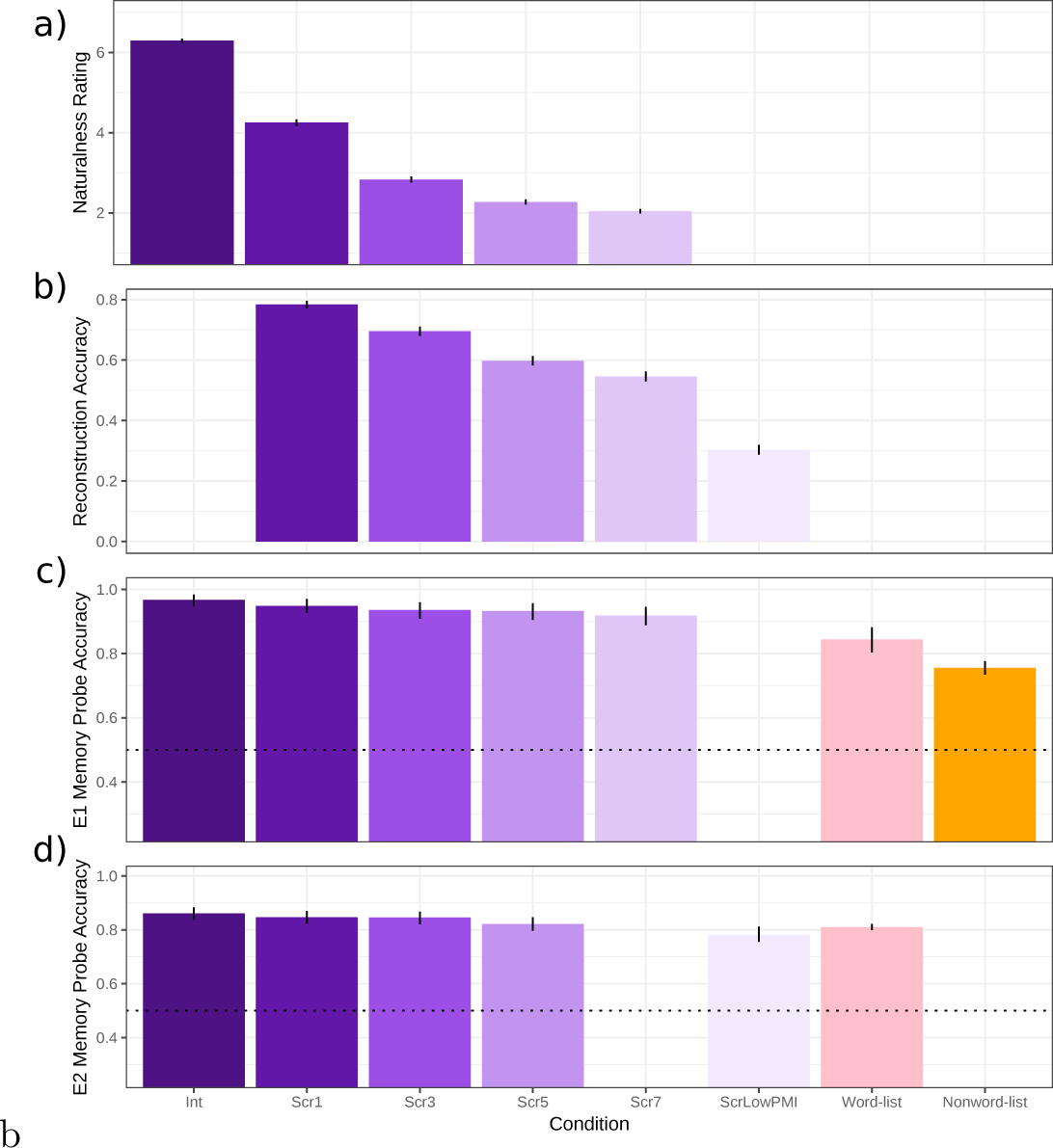
a) The average naturalness rating (higher=more natural). b) The average reconstruction accuracy. c-d) The average memory probe accuracy from Experiments 1 and 2. All error bars reflect 95% bootstrapped confidence intervals.

### Behavioral sentence reconstruction study

To assess the extent to which participants might be able to reconstruct the original sentence from its scrambled version, 180 additional participants recruited through Mechanical Turk were presented with the scrambled stimuli and asked to try to create a well-formed and meaningful sentence out of the words. As part of the instructions, several simple examples were provided. Participants were instructed that the actual stimuli would be more difficult and that they should try their best before moving on. For control purposes, we included one of the word-list conditions from Experiment 2, but we don’t analyze those data here. Similar to the rating study, the materials were distributed across experimental lists (six lists in this study, 150 trials each) following a Latin Square design. Each list contained only one version of a sentence and 25 trials of each of the six conditions (Scr1, Scr3, Scr5, Scr7, ScrLowPMI, word-list). Any given participant saw the materials from just one experimental list. Each list was seen by 30 participants. On each trial, participants were presented with a single stimulus on the screen along with a text box. Participants’ responses were automatically constrained to only include words in the stimulus; however, due to a script error, participants were allowed to use some words from the stimulus multiple times or omit words. In the analyses, we excluded all trials in which a response was not the same length as the stimuli, resulting in 17% overall data loss (Scr1: 6%, Scr3: 11%, Scr5: 14%, Scr7: 19%; ScrLowPMI: 34%). The distribution of data loss over conditions is itself a reflection of increasing reconstruction difficulty as the number of swaps increases.

Reconstruction accuracy was analyzed using a logistic mixed effect linear regression model with a fixed effect and random slopes for Condition, and random effects for Participant and Item. As in the naturalness rating study, Condition was backwards difference coded. As can be seen in Figure 4b and Table 3, every increase in degradation was associated with a significant decrease in the ability to reconstruct the sentence. This result suggests that it is unlikely that participants were able to reconstruct a full-fledged sentence-level meaning, especially given the word-by-word presentation and time demands of our task in the scanner compared to the unlimited time participants were given in the web-based reconstruction task. We return to this point in the Discussion.

**Table 3:**
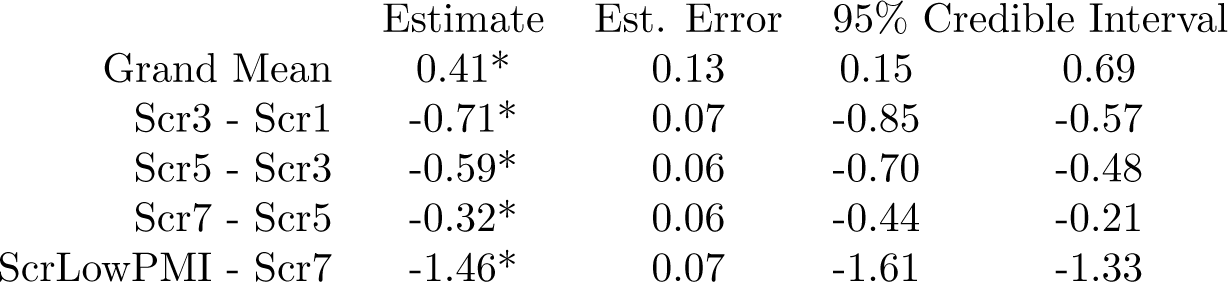
The results of a mixed effect logistic regression for the reconstruction accuracy data. * denotes significant difference.

|  | Estimate | Est. Error | 95% Credible Interval |  |
| --- | --- | --- | --- | --- |
| Grand Mean | 0.41* | 0.13 | 0.15 | 0.69 |
| Scr3 - Scr1 | -0.71* | 0.07 | -0.85 | -0.57 |
| Scr5 - Scr3 | -0.59* | 0.06 | -0.70 | -0.48 |
| Scr7 - Scr5 | -0.32* | 0.06 | -0.44 | -0.21 |
| ScrLowPMI - Scr7 | -1.46* | 0.07 | -1.61 | -1.33 |

### Discovering and characterizing brain regions sensitive to the sentence-scrambling manipulation

Given that in the behavioral naturalness rating study we found robust sensitivity to the scrambling manipulation, we asked whether any parts of the brain work harder when we process scrambled sentences. To search for brain regions sensitive to scrambling, we performed a group-constrained subject-specific (GSS) whole-brain analysis (Fedorenko et al., 2010; Julian et al., 2012). This analysis searches for spatially consistent (across individuals) patterns of activation while taking into account inter-individual variability in the precise loci of activations, which increases sensitivity relative to traditional random-effects analyses that assume voxel-wise correspondence across people (Nieto-Castañón & Fedorenko, 2012). We chose a contrast between the most scrambled condition that was shared between the two experiments (i.e., Scr5) and the Intact condition. Pooling data across experiments (n=47; for the participant who took part in both Experiments 1 and 2, we used the data from Experiment 1; for the participant who took part in Experiment 2 twice, we used the data from the first session), we took individual whole-brain activation maps for the *Scr*5 *> Int* contrast and binarized them so that voxels that show a reliable effect (significant at *p <* 0.05, uncorrected at the whole-brain level) were turned into 1’s and all other voxels were turned into 0’s. (We chose a liberal threshold for the individual activation maps to maximize our chances of detecting regions of interest; as explained below, however, the resulting regions were subsequently evaluated using statistically conservative criteria.) We overlaid these maps to create a probabilistic activation overlap map, thresholded this map to only include voxels where at least 4 of the 47 participants showed activation, and divided it into “parcels” using a watershed image parcellation algorithm (see Fedorenko et al., 2010, for details). Finally, we identified parcels that—when intersected with the individual activation maps—contained supra-threshold (i.e., significant for our contrast of interest at *p <* 0.05, uncorrected) voxels in at least half of the individual participants.

To characterize the functional profiles of scrambling-responsive regions in greater detail, in each of the regions, we estimated the BOLD response magnitude to the conditions of the two experiments. To estimate the responses to the Scr5 and Int conditions, which were used in the localizer contrast, we used an across-runs cross-validation procedure (e.g., Nieto-Castañón & Fedorenko, 2012), to ensure independence between the data used to define the fROIs and to estimate the responses (Kriegeskorte, Simmons, Bellgowan, & Baker, 2009). In particular, each parcel was intersected with each participant’s activation map for the *Scr*5 *> Int* contrast for all but the first run of the data. The voxels within the parcel were sorted—for each participant— based on their *t*-values, and the top 10% of voxels were selected as that participant’s fROI. The responses were then estimated using the left-out run’s data. The procedure was repeated iteratively leaving out each of the runs. Finally, the responses were averaged across the left-out runs to derive a single response magnitude per subject per region per condition. To estimate the responses to the other critical conditions, we used data from the Scr5 and Int conditions across all runs. Statistical tests were performed on these extracted percent BOLD signal change values.

In addition, we estimated the BOLD responses of scrambling-responsive regions to two other experiments: i) the language localizer, and ii) the spatial working memory (WM) experiment. Responses to the language localizer conditions can tell us whether the scrambling-responsive regions show a signature of the language network: i.e., stronger responses to sentences than nonword sequences. We have constrained our definition of the language-responsive regions in the critical analyses by a set of parcels derived based on activations for the language localizer contrast in a large number of individuals (as described above). Thus regions outside of this network of language-responsive regions should not show language-responsive properties. So this analysis provides a reality check of sorts. Responses to the conditions of the spatial WM task tell us whether the scrambling-responsive regions may belong to the domain-general multiple demand (MD) network, which responds robustly to this task (e.g., Fedorenko et al., 2013) and which has been generally implicated in executive functions like working memory and cognitive control (Duncan, 2010, 2013).

## Results

### Behavioral (memory probe task) data in Experiments 1 and 2

Response accuracy for each experiment was analyzed with a logistic mixed effect linear regression model with a fixed effect and random slopes for Condition, and random intercepts for Participant and Item. Condition was dummy-coded with Intact Sentences as the reference level. For both experiments, accuracy was above chance for all conditions. In Experiment 1, accuracies in the scrambled sentence conditions did not significantly differ from accuracy in the intact sentence condition; however, accuracy was significantly lower in the word-list and nonword-list conditions compared to the intact sentence condition (Figure 4c and Table 4), in line with prior work (e.g., Fedorenko et al., 2010). Similarly, in Experiment 2, accuracies in the scrambled sentence conditions did not significantly differ from accuracy in the intact sentence condition; however, accuracy was lower in the ScrLowPMI and the word-list conditions compared to the intact sentence condition (Figure 4d and Table 4). (Data and analysis code are available at OSF: https://osf.io/y28fz/.)

**Table 4:** The results of logistic mixed effect models for Experiments 1 and 2 for the memory probe data. Stimulus Type was dummy-coded with Intact sentences as the reference level. * denotes significant difference.

|  |  | Experiment 1 |  |  |  |  | Experiment 2 |  |  |
| --- | --- | --- | --- | --- | --- | --- | --- | --- | --- |
|  | Estimate | Est. Error | 95% CI |  |  | Estimate | Est. Error | 95% CI |  |
| Intact (Int) | 3.63* | 0.33 | 3.03 | 4.32 |  | 1.98* | 0.24 | 1.52 | 2.46 |
| Scr1 vs Int | 0.37 | 0.86 | -0.94 | 2.44 |  | -0.19 | 0.22 | -0.62 | 0.27 |
| Sc3 vs Int | -0.02 | 0.74 | -1.19 | 1.76 |  | 0.11 | 0.27 | -0.39 | 0.70 |
| Sc5 vs Int | -0.39 | 0.59 | -1.39 | 0.94 |  | -0.27 | 0.24 | -0.71 | 0.22 |
| Scr7 vs Int | -0.45 | 0.61 | -1.50 | 0.87 |  | — | — | — | — |
| ScrLowPMI vs Int | — | — | — | — |  | -0.64* | 0.23 | -1.07 | -0.16 |
| Words vs Int | -1.74* | 0.38 | -2.51 | -1.01 |  | -0.30* | 0.17 | -0.64 | 0.04 |
| Nonwords vs Int | -2.35* | 0.33 | -3.04 | -1.75 |  | — | — | — | — |

### fMRI data in Experiments 1 and 2

In Experiment 1, replicating much prior work (Fedorenko et al., 2010; Pallier et al., 2011), well-formed sentences elicited significantly stronger BOLD responses than the word-list and nonword-list conditions (Figure 2c, Table 5). However, degrading the sentences by introducing local word swaps did not decrease the magnitude of the language network’s response: even stimuli with seven word swaps (e.g., *their last on they overwhelmed were day farewell by messages and gifts*; Figure 1) elicited as strong a response as fully grammatical sentences (e.g., *on their last day they were overwhelmed by farewell messages and gifts*; Figure 2c, Table 5). The results also held—both qualitatively and statistically—for each language ROI separately (Figure 3 and Table 6). This pattern of similarly strong responses for the well-formed and degraded sentences suggests that inter-word dependencies are being formed even when the word order violates the rules of the language, and supports the idea that composition is the core computation implemented in the language network.

**Table 5:**
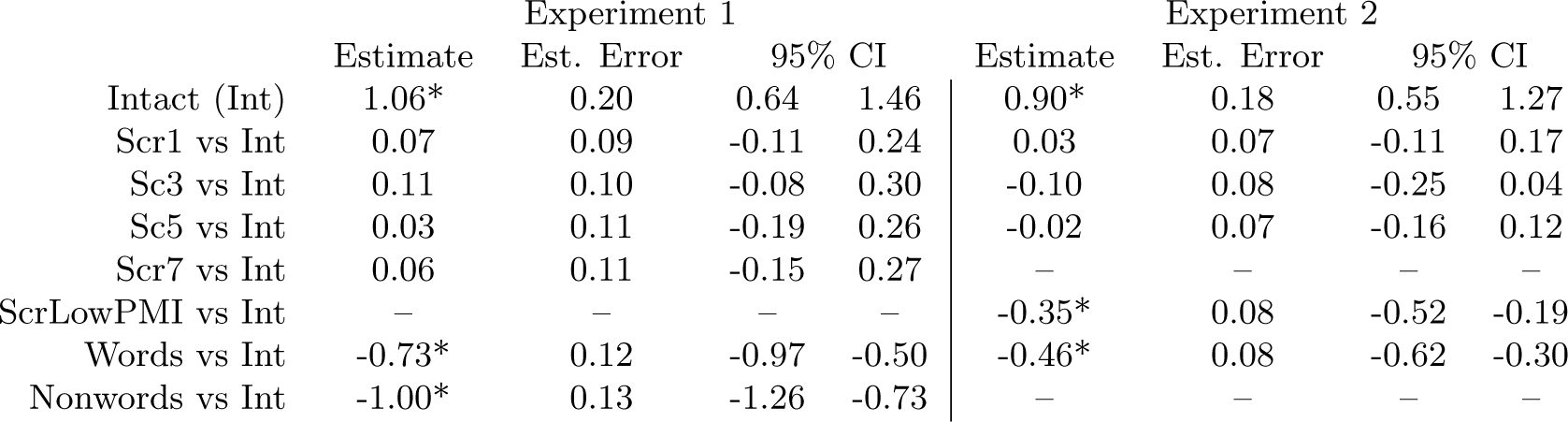
The results of mixed effect linear regressions for Experiments 1 and 2. Condition was dummy-coded with Intact sentences as the reference level. * denotes significant difference.

**Table 6:**
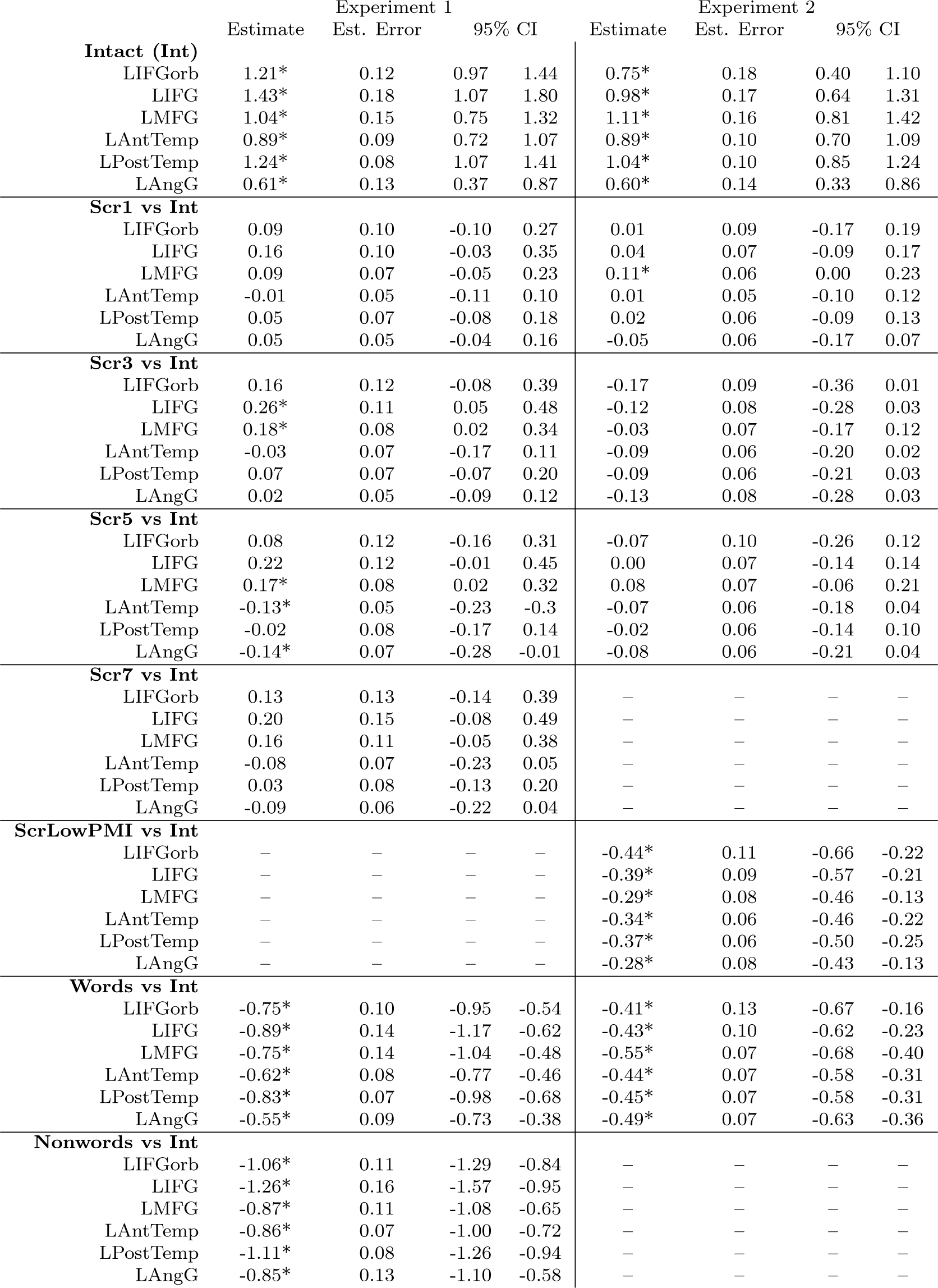
The results of mixed effect linear regressions for Experiments 1 and 2 for the six language fROIs. Condition was dummy-coded with Intact sentences as the reference level. * denotes significant difference.

In Experiment 2, we replicated the pattern observed in Experiment 1 for the intact sentences and sentences with 1, 3, or 5 local word swaps, all of which elicited similarly strong BOLD responses, all reliably higher than the control, word-list, condition (Figure 2d, Table 5). However, the ScrLowPMI condition elicited a response that was as low as that elicited by lists of unconnected words (Figure 2d, Table 5), demonstrating that combinable words have to occur in close proximity to one another for the composition mechanisms to get triggered. Again, the results held for each language ROI separately (Figure 3 and Table 6).

### Brain regions sensitive to the sentence-scrambling manipulation

In spite of eliciting as strong a BOLD response as well-formed and meaningful sentences, the scrambled sentences were rated as less acceptable behaviorally (Figure 4a and Table 2), suggesting there has to be a cost to the processing of this kind of degraded linguistic input. The whole-brain search for scrambling-sensitive areas discovered four regions, located in the middle frontal gyrus bilaterally and in the SMA (Figure 5a).

**Figure 5:**
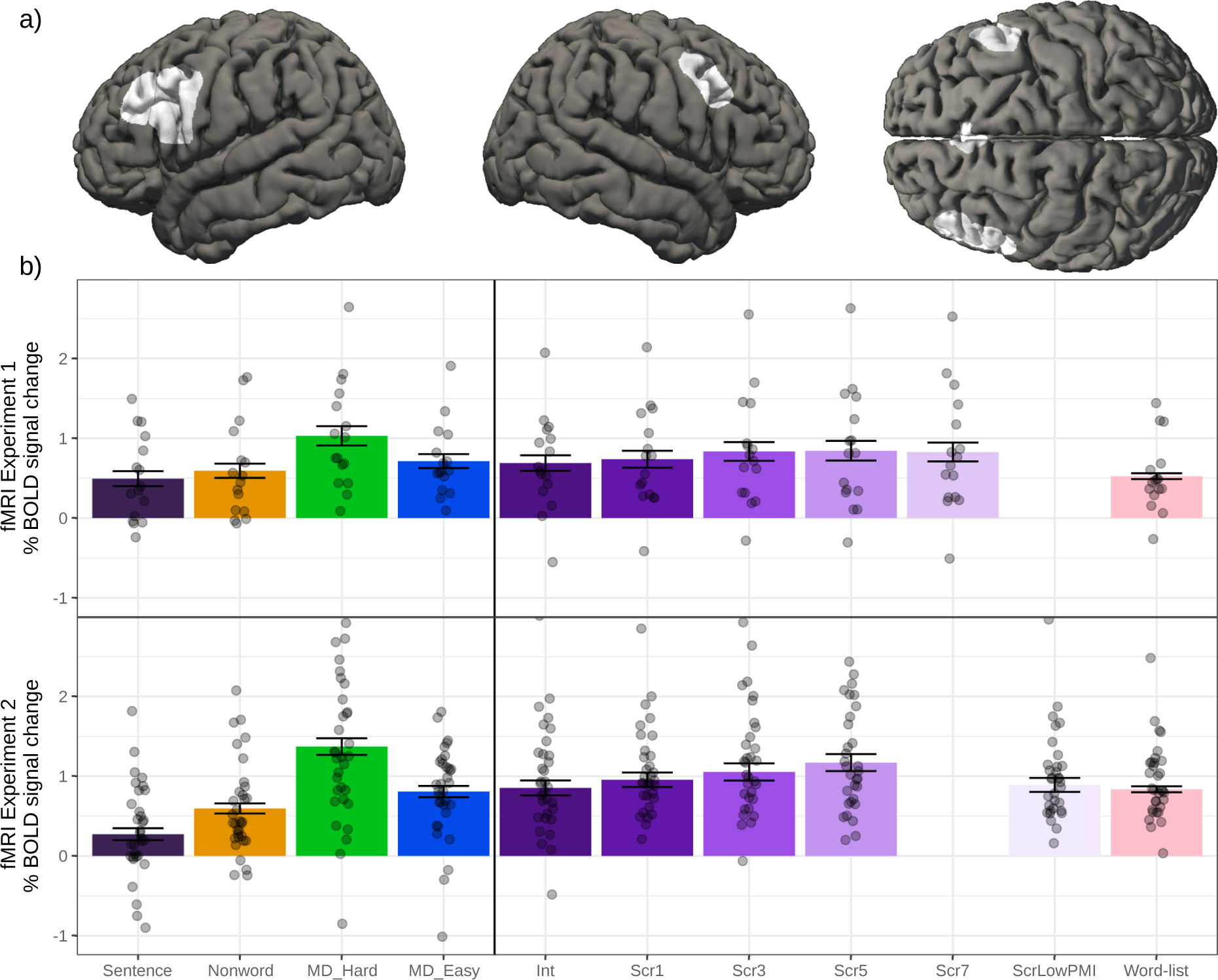
a. The parcels used to define the scrambling-responsive areas. In each participant, the top 10% of most localizer-responsive voxels within each parcel were taken as that participant’s region of interest. b. Neural responses (in % BOLD signal change relative to fixation) to the conditions of Experiments 1 (top) and 2 (bottom), as well as the language localizer and spatial WM task.

The patterns of responses observed—averaging across the fROIs—are shown in Figure 5b. Qualitatively, with respect to the conditions of the critical experiments, we found that the response increased parametrically from the Int to the Scr5 condition in both experiments. Further, in Experiment 1, the response remained high for the Scr7 condition, but in Experiment 2, the response fell off for the ScrLowPMI condition. To quantify this non-monotonic pattern, we collapsed across experiments and conducted a mixed effect linear regression with first and second order terms for Edit Distance (i.e., the number of swaps required to reconstruct the original intact sentence) as a fixed effect and random slopes, and random effects for Participant and Region of Interest. We found a small but significant increase in the BOLD response as stimuli become more scrambled, with a decrease in the ScrLowPMI condition (Table 7).

**Table 7:**
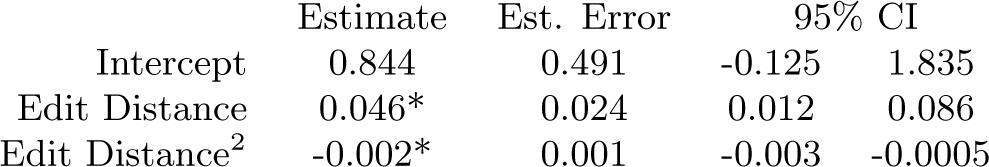
The results of mixed effect linear regression models for the scrambling-responsive regions. * denotes significant difference.

With respect to the conditions of the language localizer and the spatial WM experiment, none of the four fROIs showed a stronger response to sentences than nonword sequences (in fact, three of the four regions showed a reliably stronger response to nonword sequences than sentences, in line with Fedorenko et al. (2013); and all four fROIs showed a stronger response to the Hard than Easy condition in the spatial WM experiment. These results suggest that the scrambling-responsive fROIs fall within the domain-general MD cortex (Duncan, 2010, 2013). The parametric increase as a function of the degree of scrambling in the critical experiments is in line with robust sensitivity of the MD cortex to effort across domains (e.g., Duncan & Owen, 2000; Hugdahl et al., 2015). In particular, participants have to exert greater cognitive effort to extract meaning from the more scrambled sentences (perhaps due to greater uncertainty about how words go together, as suggested by the results of the behavioral sentence reconstruction experiment; Figure 4b). The fall-off in these fROIs for the ScrLowPMI condition—which elicited a low response in the language regions as shown in our critical analysis—is consistent with the idea that participants “give up” their attempts to derive a meaningful representation in this condition (Callicott et al., 1999; Linden et al., 2003) (cf. Wen, Mitchell, & Duncan, 2018). In particular, because participants no longer have the evidence in the input that nearby words are combinable, they stop engaging their composition mechanisms.

## Discussion

In this study, we evaluated a hypothesis that that the core linguistic computation implemented in the language-selective cortex has to do with combining words into phrases and clauses, and that this computation can take place even when the word order is not licensed by the language’s grammar. Across two fMRI experiments, we examined the processing of stimuli where the word order was degraded, via a novel parametric manipulation (varying numbers of local word swaps), making word-order-based parsing difficult or impossible, but semantic and syntactic dependencies could still be formed among nearby words. Using behavioral measures in independent groups of participants, we established robust sensitivity to the scrambling manipulation: sentences with more word swaps, and correspondingly more syntactic dependencies disrupted (Table 1), were rated as less natural (Table 2, Figure 4a), and it was more difficult to reconstruct the original sentence from the scrambled versions (Table 3, Figure 4b). However, scrambled sentences, even the conditions with a large number (5 and 7) word swaps, elicited BOLD responses in the language areas that were as strong as the response elicited by naturalistic sentences. Only when inter-word dependencies could not be formed among nearby words did the BOLD response in the language areas drop to the level of that elicited by lists of unconnected words. These results suggest that the ability to form local dependencies is necessary and sufficient for eliciting the maximal BOLD response in the language-selective brain network, where maximal is defined as the BOLD response to the preferred stimulus—well-formed and meaningful sentences. We interpret these findings as suggesting that composition is the core linguistic computation driving the neural responses in the language-selective cortex, and this computation does not depend on word order (see Bornkessel-Schlesewsky, Schlesewsky, Small, & Rauschecker, 2015, for a related proposal).

Our analyses of the experimental materials and the behavioral sentence reconstruction study help rule out two alternative explanations of these findings. One possibility is that conditions with 1, 3, 5 and even 7 word swaps (Scr1, Scr3, Scr5 and Scr7) but not the ScrLowPMI condition contained a sufficiently long well-formed and meaningful substring, and that such substrings are sufficient to elicit a BOLD response similar to that elicited by a fully well-formed sentence. To rule out this possibility, we turn to an earlier fMRI study by Pallier et al. (2011). They examined responses to 12-word long sequences that varied in their composition between a sentence, two 6-word-long substrings, three 4-word-long substrings, four 3-word-longs substrings, six 2-word-long substrings, and a list of 12 unconnected words. The BOLD response was shown to fall off as a function of the length of the substrings: a 12-word-long sentence elicited a stronger response than a sequence composed of two 6-word-long substrings, which, in turn, elicited a stronger response than a sequence composed of three 4-word-long substrings, etc. We replicate this finding in our work (Mollica et al., in prep.). The analysis of our experimental materials (Table 1) revealed that the length of the longest well-formed and meaningful substring decreases with each scrambling level and drops to 4.46 words on average for the condition with 7 word swaps. As a result, the alternative hypothesis considered here predicts a gradual fall-off in the BOLD response from the Int condition to the Scr7 condition, which is not the pattern we observe.

Another possibility is that participants were able to reconstruct the original sentence in all the scrambled conditions except for the ScrLowPMI condition. This possibility is unlikely given that in the behavioral sentence reconstruction study the ability to reconstruct the original sentence dropped off with each additional scrambling level (Figure 4b and Table 3). And this pattern was observed in spite of the fact that participants had access to the entire stimulus string and were not limited time-wise (cf. the word-by-word relatively fast presentation in the scanner). Furthermore, we model the BOLD response during the entire trial, and, by design, participants do not have access to all the words until after the last word has been presented. As a result, the similarly strong BOLD response across the Int through Scr7 conditions is unlikely due to participants successfully “unscrambling” the stimuli and processing them as such. (Of course, some local unscrambling could still take place. However, importantly, this unscrambling was, apparently, not carried out by the language areas given that there was no increase in neural response to the scrambled compared to intact stimuli in these areas. Thus, whatever computation is performed by the language areas proceeds in the same way in the intact and the scrambled conditions.)

Having ruled out these two alternatives, we argue that during the incremental processing of linguistic strings, participants form dependency relationships among words within a moving local context of a few words. This process results in the construction of phrase- and clause-level meanings. Composition is driven by the lexico-semantic and syntactic (part of speech and morphological endings) properties of the input words combined with a plausibly Bayesian inductive inference process (e.g., Steyvers, Griffiths, & Dennis, 2006). In particular, when linguistic data under-constrain interpretation, participants likely make their best guesses about the intended meaning by combining the information in the input with their prior semantic and linguistic knowledge (see e.g., Chater & Manning, 2006; Gibson et al., 2013, for applications of the general Bayesian framework to linguistic interpretation).

In addition to the consistently high BOLD response across the scrambled conditions, the behavioral data from the memory probe task performed in the scanner (Figure 4c-d and Table 4) provide indirect evidence that complex meanings were formed during the processing of all the sentence conditions except for the ScrLowPMI condition. In particular, a classic finding in the memory literature is that people’s memory for phrases and sentences is superior to their memory for lists of unconnected words (e.g., Brener, 1940; A. D. Baddeley, Hitch, & Allen, 2009), which has been attributed either to the fact that people represent sentences in terms of their meaning / gist extracted during comprehension, and that gist can later be used to regenerate the specific word-forms (e.g., Potter & Lombardi, 1990), or to the automatic engagement of long-term memory mechanisms during sentence-level comprehension, which leads to more effective binding of information within the episodic buffer (A. Baddeley, 2000; A. D. Baddeley, Allen, & Hitch, 2011). We found that participants’ performance on the memory probe task did not decline as a function of the scrambling manipulation. It only dropped in the ScrLowPMI condition. This consistently high memory probe performance can be used to indirectly infer that participants successfully formed complex meaning representations in the scrambled conditions, as they did when processing well-formed and meaningful sentences.

In the remainder of the paper, we discuss three issues that our results speak to.

### The relationship between lexico-semantic and syntactic processes

In this study, we showed that word order—one component of syntax—does not appear to affect the basic composition process carried out by the core fronto-temporal language-selective network: as long as dependencies can be formed between nearby words in linguistic strings, the composition mechanisms get engaged as they do when we process naturalistic linguistic input. Throughout the manuscript, we have talked about the composition process as encompassing both semantic composition and syntactic structure building. The relationship between the two has been treated differently across proposals in the theoretical linguistic literature. In mainstream generative grammar and formal semantics (e.g., Chomsky, 1965, 1981; Montague, 1974; B. Partee, 1975, 1995; B. B. Partee, ter Meulen, & Wall, 1990), semantic composition is considered to be a special case of syntactic composition, and syntax determines the meaning of a phrase or a clause. However, according to an alternative perspective, semantic composition can proceed (partially or fully) independently from syntactic structure building (e.g., Jackendoff & Jackendoff, 2002; Jackendoff, 2007; Culicover & Jackendoff, 2006; Culicover, Jackendoff, Jackendoff, et al., 2005; Kuperberg, 2007; Jackendoff, 2010; Jackendoff & Wittenberg, 2017). Baggio (2018) refers to this idea as “autonomous semantics”. According to his proposal, words are bound into “relational structures” based on associative, categorical, and logical relationships (see Michalon & Baggio, 2019, for evidence of computational feasibility). We are sympathetic to the latter view. We think of semantics as an independent computational system that obeys its own rules for how words are bound together during language comprehension. Of course, many of these rules have correlates in syntax, but nevertheless we conceive of semantic composition as a process that can take place independently from syntactic structure building.

However, at the implementation level, it does not appear to be the case that semantic composition and syntactic structure building are spatially separable in the brain, at least at the resolution accessible to current imaging techniques. Many have searched for and claimed to have observed a dissociation between brain regions that support (lexico-)semantic processing and those that support syntactic processing (e.g., Dapretto & Bookheimer, 1999; Embick, Marantz, Miyashita, O’Neil, & Sakai, 2000; Friederici, Opitz, & Von Cramon, 2000; Noppeney & Price, 2004; Cooke et al., 2006, inter alia). However, some of these classic findings do not appear robust to replication (Siegelman et al., 2019). And in general, taking the available evidence from cognitive neuroscience en masse, the picture that has emerged does not support a double dissociation between lexico-semantic and syntactic processes.

First, the specific regions that have been argued to support (lexico-)semantic vs. syntactic processing, and the precise construal of these regions’ contributions, differ widely across studies and proposals (e.g., Friederici, 2011, 2012; Baggio & Hagoort, 2011; Tyler et al., 2011; Bemis & Pylkkänen, 2011; Duffau, Moritz-Gasser, & Mandonnet, 2014; Ullman, 2004, 2016; Matchin & Hickok, 2019). Second, although diverse paradigms have been used across studies to probe semantic vs. syntactic processing, any given study (cf. Fedorenko, Mineroff, Siegelman, & Blank, 2018) has typically used a single paradigm, raising the possibility that the results reflect paradigm-specific differences between conditions rather than a general difference between semantic and syntactic computations. In addition, given the tight link between meaning and structure, results from some syntactic manipulations may, in fact, be due to parallel semantic composition processes. And finally, a number of neuroimaging studies have failed to observe a double dissociation between semantic and syntactic processing, reporting instead overlapping areas of activation (e.g., Keller et al., 2001; Röder, Stock, Neville, Bien, & Ræsler, 2002; Fedorenko et al., 2010; Bautista & Wilson, 2016). In particular, any brain region that shows sensitivity to syntactic processing appears to be at least as sensitive to individual word meanings and semantic composition. (There do exist brain areas—in the left anterior temporal lobe / temporal pole—that respond to word meanings, or abstract conceptual representations, according to some accounts, but not syntactic/combinatorial processing ((e.g., Schwartz et al., 2009; Schwartz, Marin, & Saffran, 1979; Patterson, Nestor, & Rogers, 2007; Visser, Jefferies, & Lambon Ralph, 2010; Mesulam et al., 2013; Lambom Ralph, Jefferies, Patterson, & Rogers, 2017)(cf. Westerlund & Pylkkänen, 2017).) In summary, it appears that syntactic processing a) is not focally carried out in a particular brain region within the language network contra some proposals (e.g., Friederici, Bahlmann, Heim, Schubotz, & Anwander, 2006; Tyler et al., 2011; Brennan et al., 2012; Berwick, Friederici, Chomsky, & Bolhuis, 2013; Matchin & Hickok, 2019), but is distributed across the left lateral frontal and temporal areas (e.g., I. Blank, Balewski, Mahowald, & Fedorenko, 2016), and b) is supported by the very same brain regions that support the processing of word meanings and semantic composition.

We would further argue that semantic composition, not syntactic structure building—to the extent that the two are separable—is primary in language comprehension and is the core operation driving the language-selective areas (see also Fedorenko et al., 2016; Pylkkänen & Brennan, in press). On the theoretical side, this argument is motivated by a key function of language—to communicate meanings (e.g., Goldberg, 2006; Jackendoff, 2011)(cf. Chomsky, Noam, et al., 2002). Abundant evidence now suggests that many properties of human languages—from the sound systems, to lexicons, to grammars—have been shaped by communicative pressures, to optimize information transfer (see Gibson et al., 2019, for review). As a result, it seems likely that our language processing mechanisms would be optimized for extracting meaning from the signal. On the empirical side, we know that meaningful sentences elicit stronger responses in the language areas than structured but meaningless stimuli, like Jabberwocky sentences or nonsensical sentences (e.g., Humphries, Binder, Medler, & Liebenthal, 2007; Fedorenko et al., 2010; Scott et al., 2017)(cf. Pallier et al., 2011), although the lack of a difference in the mean response to real vs. Jabberwocky sentences in some language areas does not appear to be replicable, and is likely driven by a between-subjects comparison in the original study (Dehaene and Pallier, p.c.)), suggesting that syntactic structure building alone cannot explain the response properties of the language areas. However, future studies should aim to further evaluate the relative importance of semantic vs. syntactic composition in language comprehension.

Our results also speak to a differential role of language statistics in syntax versus semantics. On the one hand, language statistics are relevant because humans plausibly store and continually update an implicit predictive model of linguistic forms that they use to anticipate upcoming linguistic elements during comprehension (J. Hale, 2001; Levy, 2008a; Christiansen & Chater, 2016). Indeed, a wealth of evidence demonstrates that expectations over linguistic forms affect language processing (e.g., Dell & Chang, 2014; Federmeier, 2007; Kuperberg & Jaeger, 2016; Pickering & Garrod, 2013). On the other hand, as discussed in the Introduction, language statistics are relevant because they reflect the distributional properties of objects and events in the world albeit with a bias towards objects and events that are worth encoding in and communicating through language (Griffiths, Steyvers, & Tenenbaum, 2007; Andrews, Vigliocco, & Vinson, 2009). Our work, along with a recent computational model of the N400 (Rabovsky, Hansen, & McClelland, 2018), demonstrates that the brain is sensitive to language statistics as a proxy for both world states and, perhaps more clearly, the implicit semantic dependencies in world states (e.g., which properties are likely to apply to which objects, which entities are likely engage in which actions, etc.). Keeping track of these kinds of dependencies may subsume at least some of the syntactic information. For example, Rabovsky et al. (2018) show that a model trained on semantic dependencies alone captures word order effects observed in the N400 component.

### The temporal receptive window of the language areas

An important notion has been gaining ground in the recent literature: the idea of a temporal receptive window (TRW) of a brain unit (cell, voxel, brain area) (e.g., Hasson, Yang, Vallines, Heeger, & Rubin, 2008; Lerner, Honey, Silbert, & Hasson, 2011; Overath, McDermott, Zarate, & Poeppel, 2015). A TRW is defined by Hasson and colleagues as “the length of time before a response during which sensory information may affect that response”, although the amount of information rather than time may be more relevant, especially for higher-level areas (Vagharchakian et al., 2012). What is the size of the TRW of the core language areas?

We have known for some time that discourse-level processing—connecting sentences into coherent texts— is carried out by regions outside of the fronto-temporal language network (e.g., Ferstl, Neumann, Bogler, & Von Cramon, 2008; Ferstl & von Cramon, 2001; Kuperberg, Caplan, Sitnikova, Eddy, & Holcomb, 2006; Lerner et al., 2011)(see Jacoby & Fedorenko, 2018, for evidence of insensitivity to discourse-level processing in the functionally defined languages areas of the core fronto-temporal network). For example, Lerner et al. (2011) presented participants with an auditory story as well as the same story scrambled at different grains of information (at the paragraph level, at the sentence level, and at the word level). In a whole-brain voxelwise analysis of inter-subject correlations (Hasson et al., 2008), which can be used to draw inferences about the size of the TRW of a voxel, they found that a) brain areas sensitive to paragraph-level structure and above resemble the Default Mode network (e.g., Buckner, Andrews-Hanna, & Schacter, 2008) or the network that supports social cognition (e.g., Saxe & Kanwisher, 2003), and b) brain areas sensitive to word and sentence-level processing (but not to structure above the sentence level) resemble the core language network. The inter-subject correlations were higher for the sentence-scrambled condition than the word-scrambled condition (see also I. A. Blank & Fedorenko, 2019) but where exactly between a single word and a sentence does the TRW of the language areas fall?

Pallier et al. (2011)’s study discussed above showed that the response in the language network appears to increase gradually from same-length sequences composed of single words to 2-word phrases, to 3-word phrases, to 4-words phrases, to 6-word phrases, with an additional, albeit smaller increase for full sentences. Our results suggest that when combinable words are separated by 8 words (as previously adjacent content words are in the ScrLowPMI condition; average separation is 8.33 words), resulting in low average local PMI, composition does not take place as evidence by a low response in the language areas. The TRW of the language areas therefore appears to be in the 5-7 word range. As alluded to in the Introduction, this relatively local linguistic processing is likely driven by the statistical properties of natural language, where most semantic/syntactic dependencies are local (e.g., Futrell et al., 2015), and PMI falls off quite sharply as a function of inter-word distance (e.g., Lin & Tegmark, 2017). We can further speculate that linguistic chunks of this size are sufficient to express clause-level meanings, where clauses describe events – salient and meaningful semantic units in our experience with the world (e.g., Zacks & Tversky, 2001). Of course, we can detect and process syntactic and anaphoric dependencies that span much longer windows than 6 words, and these types of non-local dependencies have been extensively investigated in the psycholinguistic literature (e.g., Yngve, 1960; Miller & Chomsky, 1963; Lewis & Jones, 1996; Gibson, 1998, 2000, inter alia). How exactly the processing of such dependencies is carried out in the brain remains debated, in part because the most commonly used method in cognitive neuroscience (fMRI) lacks the temporal resolution needed to track the dynamics of dependency formation. We don’t take our results as inconsistent with the human ability to process non-local dependencies; instead, we take them to suggest that our language processing mechanisms may be optimized for dealing with particular-size packages of linguistic information.

### Sensitivity of domain-general executive mechanisms to the scrambling manipulation

Although the language-selective regions’ BOLD responses were robust to the scrambling manipulation, in a behavioral rating study, more scrambled sentences elicited lower naturalness ratings (Figure 4a and Table 2), suggesting that such sentences should incur a greater processing cost. What cognitive and neural mechanisms handle this extra cost? We found a number of brain regions that appear to fall within the domain-general multiple demand (MD) network (Duncan, 2010, 2010), which has been implicated broadly in goal-directed behavior and linked to executive functions, like working memory and cognitive control. These regions expended more energy when participants processed sentences with scrambled word orders compared to intact sentences. The level of BOLD response increased as the degree of scrambling increased, until participants were no longer able to form local semantic dependencies (as evidenced by a drop in the BOLD response in the language network), which occurred in the ScrLowPMI condition. These results suggest that the cost associated with the processing of scrambled sentences is carried by domain-general executive regions that support diverse demanding tasks across domains (e.g., Duncan & Owen, 2000; Hugdahl et al., 2015).

The importance and the precise role of the MD network in language comprehension remains debated (e.g., Wright, Randall, Marslen-Wilson, & Tyler, 2011; I. Blank et al., 2014; Campbell & Tyler, 2018; Diachek, Blank, Siegelman, & Fedorenko, 2019). A number of prior studies have reported activation in the MD areas during the processing of acoustically degraded speech (e.g., Peelle, 2018) or sentences with syntactic errors (e.g., Kuperberg et al., 2003), suggesting that the MD network may be important for coping with signal corruption, perhaps performing specific operations aimed at “repairing” the input. However, other studies have reported MD activity during conditions that do not involve corrupted input, both in the domain of language (e.g., Whitney, Kirk, O’Sullivan, Lambon Ralph, & Jefferies, 2012; Hoffman, Loginova, & Russell, 2018), and for many non-linguistic tasks (e.g., Duncan & Owen, 2000; Crittenden & Duncan, 2012; Fedorenko et al., 2013; Hugdahl et al., 2015), suggesting perhaps that the contribution is more general in nature (e.g., providing more attentional or working memory resources). At this time, it is difficult to put forward mechanistic-level accounts of the MD networks’ contribution to processing noisy linguistic input.

## Conclusion

To conclude, we have provided evidence that constructing complex meanings appears to be the core linguistic computation implemented in the language-selective fronto-temporal network: as long as that computation is engaged (as determined by the combination of input properties and a plausibly Bayesian inference process), language brain areas are as active as when they process their preferred stimulus – well-formed meaningful sentences. Moreover, combinable words have to occur in close proximity to one another for the composition mechanisms to get triggered. Many important questions about linguistic composition remain. For example, how strongly is composition driven by our prior experience with particular words vs. the underlying concepts? Is the span over which high mutual information is detected and affects composition determined by language statistics or by our general memory limitations? And is it similar between the visual and auditory modalities? How exactly do bottom-up lexico-semantic and syntactic cues trade off with top-down inferential processes that take into account our knowledge of language and the world? And how are we able to quickly re-map our world-knowledge priors when we process fictional or otherwise implausible scenarios (e.g., Nieuwland & Van Berkum, 2006)? In spite of all these open questions, current work brings us one step closer to a mechanistic-level account of the computations that the language network plausibly supports.

